# Genome-wide analysis of retinal transcriptome reveals common genetic network underlying perception of contrast and optical defocus detection

**DOI:** 10.1101/2020.09.16.300244

**Authors:** Tatiana V. Tkatchenko, Andrei V. Tkatchenko

**Author notes:** **Correspondence to**: Andrei V. Tkatchenko, Edward S. Harkness Eye Institute, Research Annex Room 415, 635 W. 165th Street, New York, NY 10032, USA.

## Abstract

Refractive eye development is regulated by optical defocus in a process of emmetropization. Excessive exposure to negative optical defocus often leads to the development of myopia. However, it is still largely unknown how optical defocus is detected by the retina. Here, we used genome-wide RNA-sequencing (RNA-seq) to conduct analysis of the retinal genetic networks underlying contrast perception and refractive eye development. We report that the genetic network subserving contrast perception plays an important role in optical defocus detection and emmetropization. Our results demonstrate an interaction between contrast perception, the retinal circadian clock pathway and the signaling pathway underlying optical defocus detection. We also observe that the relative majority of genes causing human myopia are involved in the processing of optical defocus. Together, our results support the hypothesis that optical defocus is perceived by the retina using contrast as a proxy and provide new insights into molecular signaling underlying refractive eye development.

## INTRODUCTION

Refractive eye development is controlled by both environmental and genetic factors, which determine the optical geometry of the eye and its refractive state by a process called emmetropization (1–8). Various environmental factors influence refractive eye development (8–11); however, the leading environmental factor driving emmetropization is optical defocus (8, 12). The eye is very sensitive to the sign of optical defocus and can compensate for imposed defocus very accurately by modulating the growth of the posterior segment of the eye via a developmental mechanism called bidirectional emmetropization by the sign of optical defocus (BESOD), whereby negative optical defocus stimulates eye growth and positive optical defocus suppresses it (12–18). BESOD uses optical defocus to match the eye’s axial length to its optical power and can produce either sharp vision (if the eye is exposed to a normal visual environment), myopia (if the eye is exposed to negative optical defocus) or hyperopia (if the eye is exposed to positive optical defocus) (8). Importantly, animal studies suggest that the process of emmetropization is controlled locally by the retina (19–24). Excessive exposure to negative optical defocus associated with nearwork leads to the development of myopia (nearsightedness), which is the most common ocular disorder in the world, manifesting in children of school age as blurred distance vision (13,14,16–18,25–33).

Although the role of optical defocus in refractive eye development is a well-established fact, it is still largely unknown how optical defocus is perceived by the retina. Several lines of evidence suggest that the absolute level of visual acuity does not play a significant role in optical defocus detection because a variety of species with different visual acuity ranging from 1.3 cpd in fish (34), 0.6-1.4 cpd in mice (35–37), 2.4 cpd in tree shrews (38, 39), 2.7 cpd in guinea pigs (40, 41), 5 cpd in cats (42, 43), 5 cpd chickens (44, 45) and 44 cpd in rhesus monkeys and humans (46–48) undergo emmetropization and can compensate for imposed optical defocus (13-17,49-53). Moreover, it was demonstrated that accommodation and emmetropization are driven by low spatial frequencies even in species with high visual acuity (54, 55); and the peripheral retina, which has much lower visual acuity than the central retina (56–61), plays an important role in defocus detection and emmetropization (8,22,24,62). Conversely, perception of contrast is emerging as an important cue for both defocus-driven accommodation and emmetropization (54,63,64). The possible role of contrast perception in emmetropization is highlighted by the fact that species with different visual acuity have similar contrast sensitivity at spatial frequencies found to be critical for emmetropization (65). Optical defocus leads to a proportional degradation of contrast at the luminance edges of the images projected onto the retina, as revealed by the analysis of the effect of defocus on the eye’s contrast sensitivity (66–69). The retina, as revealed by *in vitro* recordings from ganglion cells, exhibits the highest sensitivity to contrast at low spatial frequencies, consistent with what is found for the dependency of accommodation and emmetropization on low spatial frequencies (45). Ganglion cells’ firing rate increases proportionally with an increase in both optical focus and contrast, suggesting that contrast may be used by the retina as a proxy to optical defocus (45). Interestingly, vision across the animal kingdom is tuned to contrast and not to visual acuity (65,70,71). Several species have been shown to possess high contrast sensitivity and undergo emmetropization despite low visual acuity (65,72–75). Retinae of many species adapted asymmetric photoreceptor distribution which maximizes contrast perception across visual space in expense of other important visual functions such as visual acuity and color perception (70,71,76–81).

Accommodation and emmetropization appear to be driven by both luminance contrast and longitudinal chromatic aberrations, whereby the retina uses the contrast of the luminance edges to determine focal plane and color contrast to identify the sign of defocus (54,63,64,82). Detection of luminance contrast and color contrast in the retina is organized as ON-center/OFF-surround and OFF-center/ON-surround receptive fields, which divide retinal pathways into ON and OFF channels respectively (83, 84). The importance of these contrast processing retinal channels for refractive eye development was demonstrated by experiments in knockout mice with ON-pathway mutations and in humans (85–88). Anatomically, center-surround receptive fields are comprised of photoreceptors, bipolar cells, horizontal cells and amacrine cells, where horizontal and amacrine cells provide lateral inhibition which is critical for the generation of surround and contrast perception (83, 84). Interestingly, it was shown that amacrine cells play a critical role in refractive eye development (89–99) and that loss of amacrine cells negatively affects contrast sensitivity (100).

Cronin-Golomb et al. (101) found that genetic factors have a significant contribution to contrast sensitivity in humans. The retina converts information about optical defocus into molecular signals, which are transmitted across the back of the eye via a multilayered signaling cascade encoded by an elaborate genetic network (12,102–115). It is critical to characterize the genetic networks and signaling pathways underlying contrast perception and determine their role in refractive eye development.

Here, we systematically analyzed genetic networks and signaling pathways underlying contrast perception and refractive eye development in a panel of highly genetically diverse mice, which have different baseline refractive errors, different susceptibility to form-deprivation myopia and different levels of contrast sensitivity, and determined the contribution of the genetic network subserving contrast perception to baseline refractive development and optical-defocus-driven eye emmetropization.

## RESULTS

### Contrast sensitivity and susceptibility to form-deprivation myopia exhibit strong interdependence in mice

The mechanism of defocus perception by the retina remains poorly understood. To investigate the role of contrast in defocus perception and refractive eye development, we analyzed the relationship between contrast sensitivity, baseline refractive error, and susceptibility to form-deprivation myopia in eight genetically diverse strains of mice comprising collaborative cross, i.e., 129S1/svlmj, A/J, C57BL/6J, CAST/EiJ, NOD/ShiLtJ, NZO/HlLtJ, PWK/PhJ, and WSB/EiJ mice.

Analysis of baseline refractive errors and susceptibility to form-deprivation myopia in these mice (Figure 1A and B; Supplementary Tables S1 and S2) revealed that although both parameters were clearly influenced by the differences in genetic background between the strains (ANOVA_refractive error_, F(7, 145) = 429.8, P < 0.00001; ANOVA_myopia_, F(7, 48) = 9.8, P < 0.00001) and both baseline refractive error and susceptibility to form-deprivation myopia were inherited as quantitative traits, correlation between baseline refractive error and susceptibility to myopia was weak (r = 0.2686, P = 0.0305). Therefore, we then analyzed contrast sensitivity in all eight strains of mice (Figure 1C; Supplementary Table S3) and found that genetic background strongly influenced contrast sensitivity (ANOVA_contrast_, F(7, 57) = 1837.7, P < 0.00001). Moreover, contrast sensitivity was also inherited as a quantitative trait. Linear regression analysis showed that the correlation between contrast sensitivity and baseline refractive error was not statistically significant (r = 0.1058, P = 0.4014) (Figure 1D), whereas contrast sensitivity and form-deprivation myopia exhibited a positive statistically significant correlation (r = 0.5723, P < 0.0001) (Figure 1E). Mouse strain-specific contrast sensitivity profiles revealed largely consistent differences between the strains at all frequencies (Figure 1F; Supplementary Table S3), with maximum contrast sensitivity recorded at 0.064 cpd. Therefore, contrast sensitivity at 0.064 cpd was used for all further analyses.

**Figure 1.**
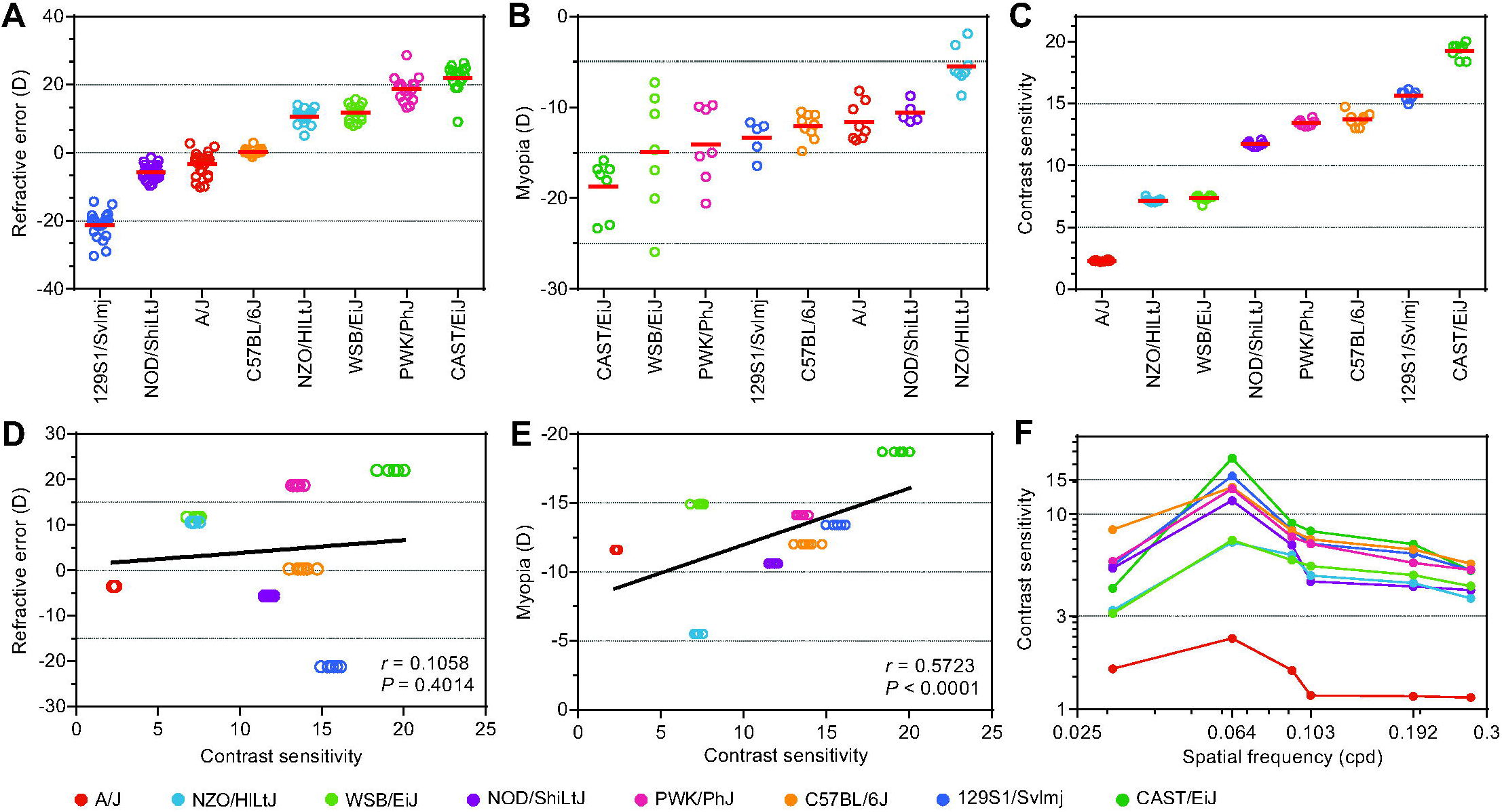
Contrast sensitivity strongly correlates with susceptibility to form-deprivation myopia in mice. (A) Baseline refractive errors range from highly myopic to highly hyperopic in mice depending on the genetic background. Horizontal red lines show mean refractive errors for each strain, while each dot corresponds to mean refractive errors of individual animals. (B) Susceptibility to form-deprivation myopia in mice is determined by genetic background. Horizontal red lines identify means of induced myopia for each strain, while each dot represents a mean interocular difference between the treated eye and contralateral control eye for individual animals. (C) Contrast sensitivity in mice is determined by genetic background. Horizontal red lines identify means of contrast sensitivity for each strain, while each dot represents a mean contrast sensitivity for individual animals. (D) Baseline refractive error does not correlate with sensitivity to contrast. Linear regression showing lack of correlation between baseline refractive error and contrast sensitivity. r, Pearson’s correlation coefficient; P, Pearson’s correlation significance. (E) Susceptibility to form-deprivation myopia correlates with sensitivity to contrast. Linear regression showing correlation between form-deprivation myopia and contrast sensitivity. r, Pearson’s correlation coefficient; P, Pearson’s correlation significance. (F) Contrast sensitivity profiles in eight genetically different strains of mice. Contrast sensitivity was measured at 0.031, 0.064, 0.092, 0.103, 0.192, and 0.272 cycles/degree (cpd). Colors identify different strains of mice as shown at the bottom.

Collectively, these data suggest that processing of contrast by the retina plays an important role in optical defocus detection, visually regulated eye growth and susceptibility to form-deprivation myopia. On the other hand, contrast sensitivity does not seem to play any substantial role in baseline refractive eye development.

### Contrast sensitivity, baseline refractive eye development, and susceptibility to form-deprivation myopia are controlled by diverse sets of genes and signaling pathways

Considering that our data suggested that contrast sensitivity exhibited strong correlation with susceptibility to form-deprivation myopia and very weak correlation with baseline refractive error, we then set out to investigate whether these observations would be reflected at the level of molecular signaling underlying contrast perception, baseline refractive eye development, and the development of form-deprivation myopia. To gain insight into the molecular signaling cascades involved in the regulation of these processes, we performed a genome-wide gene expression profiling in the retina of the eight mouse strains using massively parallel RNA-sequencing (RNA-seq).

We found that expression of at least 2,050 genes (Q-value < 0.01) correlated with baseline refractive error (Figure 2A, Supplementary Table S4). These genes were organized in two clusters (Figure 2A). The expression of 718 genes in the first cluster positively correlated with baseline refractive error (i.e., expression was increased in hyperopic animals and decreased in myopic animals), while the expression of 1,332 genes in the second cluster negatively correlated with baseline refractive error (i.e., expression was decreased in hyperopic animals and increased in myopic animals). Analysis of the gene ontology categories associated with biological processes revealed 88 biological processes, which were involved in baseline refractive eye development (Figure 2B, Supplementary Table S7). Among these processes, several were linked to visual perception, synaptic transmission, cell-cell communication, retina development, and DNA methylation, which suggests that these processes play an important role in baseline refractive eye development. Gene ontology data were complemented by the analysis of canonical signaling pathways, which revealed that signaling pathways related to β protein kinase A signaling, dopamine receptor signaling, HIPPO signaling, mTOR signaling, phototransduction pathway, axonal guidance signaling, synaptic long term potentiation, tight junction signaling, and DNA methylation signaling, among others, are involved in baseline refractive eye development (Figure 2C, Supplementary Table S10).

**Figure 2.**
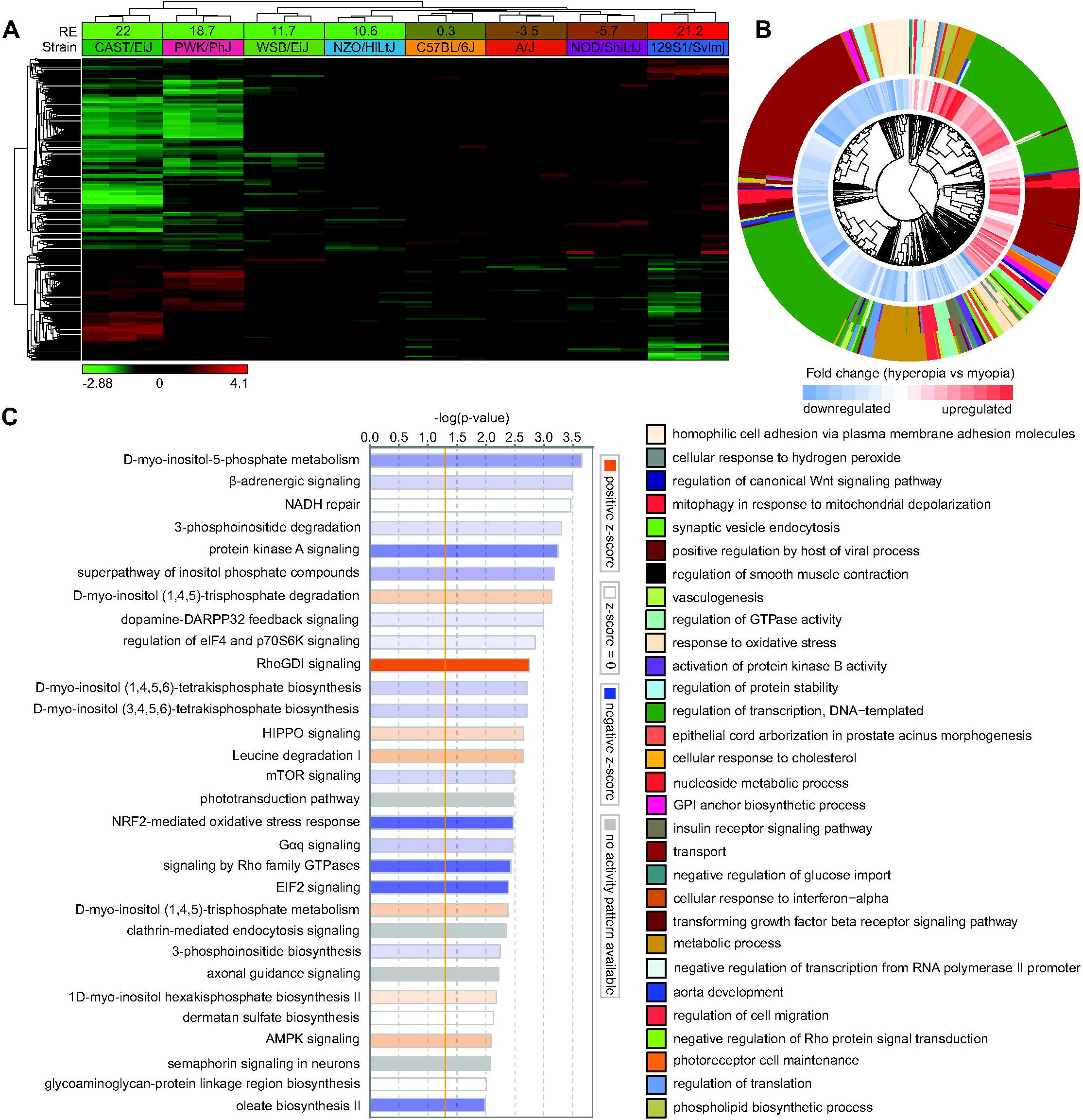
Genetic network underlying baseline refractive eye development controls diverse biological functions and signaling pathways. (A) Expression of 2,050 genes (Q-value < 0.01) is influenced by genetic background and correlates with baseline refractive errors. Hierarchical clustering shows that differential genes are organized in two clusters: one (top) cluster exhibits increased expression in the myopic mice, and the second (bottom) cluster shows increased expression in the hyperopic mice. (B) Top 30 biological processes affected by the genes involved in baseline refractive eye development. Outer circle of the hierarchical clustering diagram shows hierarchical clusters of biological processes (identified by different colors); inner circle shows clusters of the associated genes up- or down-regulated in hyperopic mice versus myopic mice. (C) Top 30 canonical pathways affected by the genes associated with baseline refractive eye development. Horizontal yellow line indicates P = 0.05. Z-scores show activation or suppression of the corresponding pathways in animals with hyperopia versus animals with myopia.

Analysis of the relationship between gene expression and susceptibility to form-deprivation myopia uncovered that expression of at least 1,347 genes (Q-value < 0.01) correlated with susceptibility to form-deprivation myopia (Figure 3A, Supplementary Table S5), including 455 genes whose expression positively correlated with susceptibility to form-deprivation myopia (i.e., expression was increased in animals with high susceptibility to form-deprivation myopia and decreased in animals with low susceptibility to form-deprivation myopia) and 892 genes which exhibited a negative correlation with susceptibility to myopia (i.e., expression was decreased in animals with high susceptibility to form-deprivation myopia and increased in animals with low susceptibility to form-deprivation myopia). Gene ontology analysis revealed potential involvement of the biological processes related to axonogenesis, transport, fatty acid oxidation, aging, insulin secretion, cell proliferation, cell-cell adhesion, and cellular response to hypoxia, among others (Figure 3B, Supplementary Table S8). Furthermore, analysis of canonical signaling pathways pointed to the important role of G2/M DNA damage checkpoint regulation, iron homeostasis signaling pathway, tight junction signaling, sirtuin signaling pathway, β adrenergic and α renergic signaling, HIPPO signaling, mTOR signaling, amyotrophic lateral sclerosis signaling, axonal guidance signaling, and growth hormone signaling, among others (Figure 3C, Supplementary Table S11).

**Figure 3.**
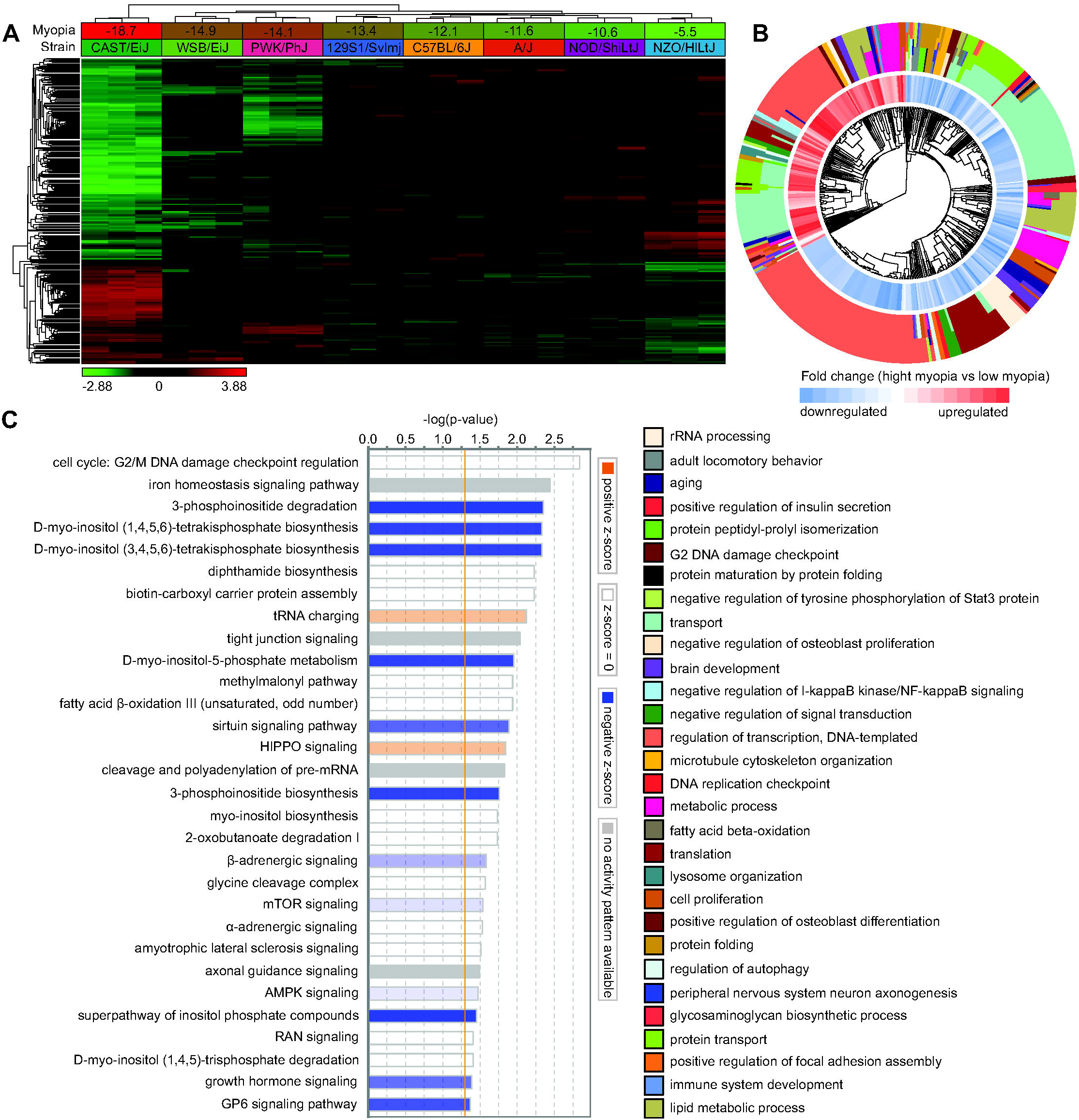
Genetic network subserving susceptibility to form-deprivation myopia controls diverse biological functions and signaling pathways. (A) Expression of 1,347 genes (Q-value < 0.01) is influenced by genetic background and correlates with susceptibility to form-deprivation myopia. Hierarchical clustering shows that differential genes are organized in two clusters: one (top) cluster exhibits increased expression in mice with low susceptibility to myopia, and the second (bottom) cluster shows increased expression in mice with high susceptibility to myopia. (B) Top 30 biological processes affected by the genes involved in optical defocus detection and the development of form-deprivation myopia. Outer circle of the hierarchical clustering diagram shows hierarchical clusters of biological processes (identified by different colors); inner circle shows clusters of the associated genes up- or down-regulated in mice with high susceptibility to myopia versus mice with low susceptibility to myopia. (C) Top 30 canonical pathways affected by the genes associated with optical defocus detection and the development of form-deprivation myopia. Horizontal yellow line indicates P = 0.05. Z-scores show activation or suppression of the corresponding pathways in animals with high susceptibility to form-deprivation myopia versus animals with low susceptibility to form-deprivation myopia.

We found that regulation of contrast sensitivity in mice is associated with differential expression of at least 1,024 genes in the retina (Q-value < 0.01) (Figure 4A, Supplementary Table S6). Expression of 489 of these genes was positively correlated with contrast sensitivity (i.e., expression was increased in animals with high contrast sensitivity and reduced in animals with low contrast sensitivity), whereas expression of 535 of these genes was negatively correlated with contrast sensitivity (i.e., expression was decreased in animals with high contrast sensitivity and increased in animals with low contrast sensitivity). Interestingly, contrast sensitivity was found to be strongly associated with biological processes related to rhythmic process, entrainment of circadian clock by photoperiod, circadian regulation of gene expression, regulation of circadian rhythm, detection of light stimulus involved in visual perception, visual perception, receptor-mediated endocytosis, synapse assembly, cell adhesion, positive regulation of catenin import into nucleus, and RNA splicing (Figure 4B, Supplementary Table S9). At the level of canonical signaling pathways, we found a strong association of contrast sensitivity with CD27 signaling, leucine degradation, senescence pathway, phototransduction pathway, melatonin signaling, synaptic long-term depression, relaxin signaling, PPARα HIPPO signaling, and DNA methylation signaling, among other pathways (Figure 4C, Supplementary Table S12).

**Figure 4.**
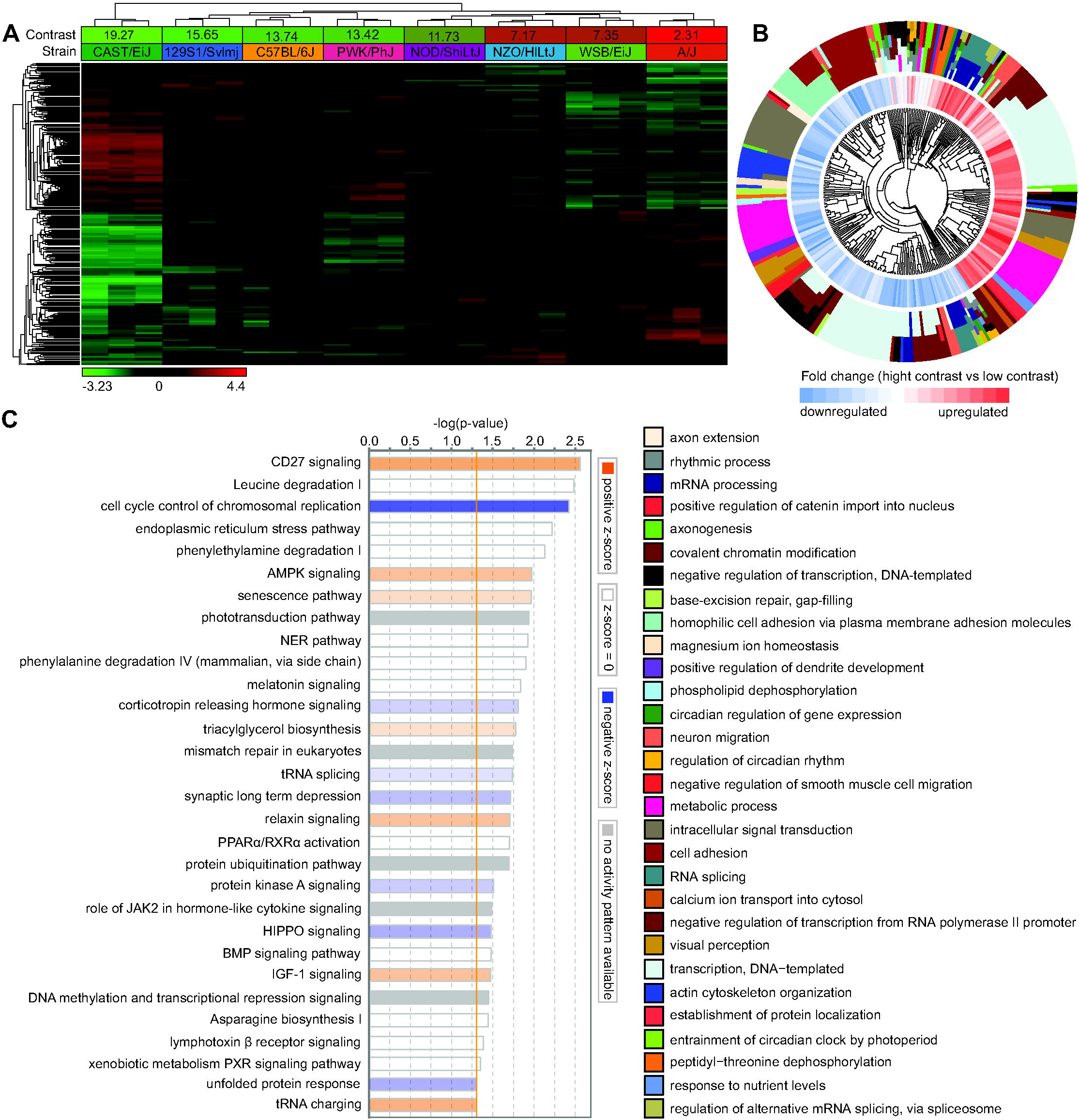
Genetic network underlying perception of contrast controls diverse biological functions and signaling pathways. (A) Expression of 1,024 genes (Q-value < 0.01) is influenced by genetic background and correlates with sensitivity to contrast. Hierarchical clustering shows that differential genes are organized in two clusters: one (top) cluster exhibits increased expression in mice with high contrast sensitivity, and the second (bottom) cluster shows increased expression in mice with low contrast sensitivity. (B) Top 30 biological processes affected by the genes involved in contrast perception. Outer circle of the hierarchical clustering diagram shows hierarchical clusters of biological processes (identified by different colors); inner circle shows clusters of the associated genes up- or down-regulated in mice with high contrast sensitivity versus mice with low contrast sensitivity. (C) Top 30 canonical pathways affected by the genes associated with contrast perception. Horizontal yellow line indicates P = 0.05. Z-scores show activation or suppression of the corresponding pathways in animals with high contrast sensitivity versus animals with low contrast sensitivity.

Taken together, these data highlight the complexity and diversity of biological processes and signaling pathways involved in refractive eye development and suggest that the correlation between contrast perception and susceptibility to form-deprivation myopia may be explained by the common signaling pathways underlying both processes. The relationship between contrast sensitivity and baseline refractive development appears to be less pronounced.

### Comparison of transcriptomes underlying contrast sensitivity and baseline refractive eye development reveals limited contribution of the genetic network regulating contrast perception to baseline refractive eye development

To elucidate the relationship between contrast perception and baseline refractive eye development, we analyzed the overlap between genetic networks, biological processes, and signaling pathways underlying contrast sensitivity and baseline refractive development. We found that less than 13% of genes involved in the regulation of contrast sensitivity (136 genes) exhibited a correlation with baseline refractive error (OR = 1.9, P = 1.05 × 10^−12^) (Figure 5A, Supplementary Table S13). These genes were involved in 11 biological processes shown in Figure 5B (Supplementary Table S15), which implicate DNA and histone methylation, as well as cell adhesion and dendrite development. Importantly, both increased contrast sensitivity and hyperopic refractive errors were associated with suppression of many of these processes. Analysis of canonical signaling pathways affected by the genes whose expression correlates with both contrast sensitivity and baseline refractive error revealed that these two processes are controlled by largely distinct pathways (Figure 5C, Supplementary Tables S10 and S12). Nevertheless, several canonical pathways involved in the regulation of contrast perception were also implicated in the regulation of baseline refractive eye development, including protein kinase A signaling, HIPPO signaling, leucine degradation, phototransduction pathway, AMPK signaling, senescence pathway, melatonin signaling, tRNA splicing, synaptic long-term /RXRα activation, xenobiotic metabolism PXR signaling pathway, unfolded protein response, and DNA methylation and transcriptional repression signaling. Thus, although the overlap between the genetic network underlying contrast perception and that underlying baseline refractive development is statistically significant, the cumulative evidence suggests that the overall impact of the genetic network subserving contrast perception on baseline refractive eye development appears to be relatively low.

**Figure 5.**
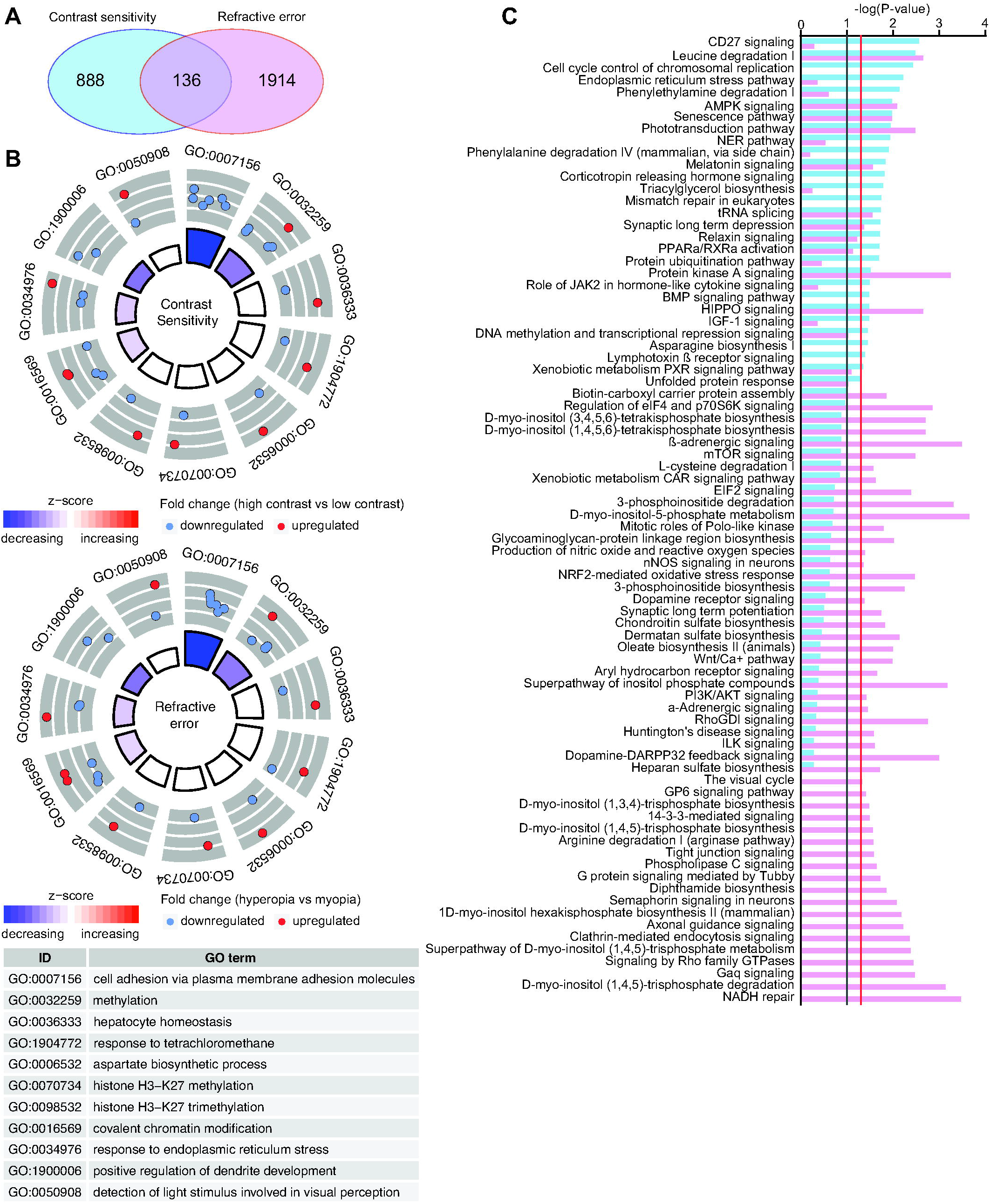
Genetic network underlying contrast perception has limited contribution to baseline refractive eye development. (A) Venn diagram showing overlap between genes underlying contrast sensitivity and genes regulating baseline refractive development. (B) Top 11 biological processes affected by the genes associated with both contrast perception (top panel) and baseline refractive development in mice (bottom panel). Outer circle shows gene ontology IDs for the biological processes; middle circle shows up- or down-regulated genes in animals with high contrast sensitivity versus animals with low contrast sensitivity (top panel), or in mice with hyperopia versus mice with myopia (bottom panel); inner circle shows activation or suppression of the corresponding biological processes, while the size of the sector corresponds to statistical significance (larger sectors correspond to smaller P-values). (C) Comparison of canonical signaling pathways involved in contrast perception and baseline refractive eye development. Vertical red line indicates P = 0.05. Vertical grey line indicates P = 0.1. Colors identify pathways associated with either contrast perception or baseline refractive development and correspond to the colors in the Venn diagrams (A).

### Comparison of transcriptomes underlying contrast sensitivity and the development of form-deprivation myopia reveals significant contribution of the genetic network regulating contrast perception to optical defocus detection and emmetropization

Considering that our data suggested that there was statistically significant correlation between contrast sensitivity and susceptibility to form-deprivation myopia, we analyzed the overlap between genes, biological processes, and canonical signaling pathways associated with contrast perception and the development of form-deprivation myopia. We discovered that more than 30% of genes (315 genes) whose expression correlates with contrast sensitivity are also involved in the regulation of susceptibility to form-deprivation myopia (Figure 6A, Supplementary Table S14). The overlap between these two gene sets was highly significant (OR = 6.6, P = 6.61 × 10^− 175^). These genes were associated with 21 biological processes, implicating regulation of circadian rhythm, positive regulation of focal adhesion assembly, cellular response to hypoxia, axon extension, and DNA methylation (Figure 6B, Supplementary Table S16). Importantly, the processes, which were activated in the animals with high contrast sensitivity, were also activated in the animals with high susceptibility to form-deprivation myopia, while the processes, which were suppressed in the animals with high contrast sensitivity, were also suppressed in the animals with high susceptibility to form-deprivation myopia. Analysis of the canonical signaling pathways encoded by the genes associated with contrast perception and susceptibility to form-deprivation myopia revealed that there was a substantial overlap between these two processes at the level of canonical signaling pathways (Figure 6C, Supplementary Tables S11 and S12). Approximately 75% of pathways linked to form-deprivation myopia development (27 out of 36) were also associated with contrast perception, including tRNA charging, HIPPO signaling, AMPK signaling, NER pathway, IGF-1 signaling, protein kinase A signaling, role of JAK2 in hormone-like cytokine signaling, relaxin signaling, and PPARα these data suggest that the genetic network regulating contrast perception significantly contributes to optical defocus detection and visually guided eye emmetropization.

**Figure 6.**
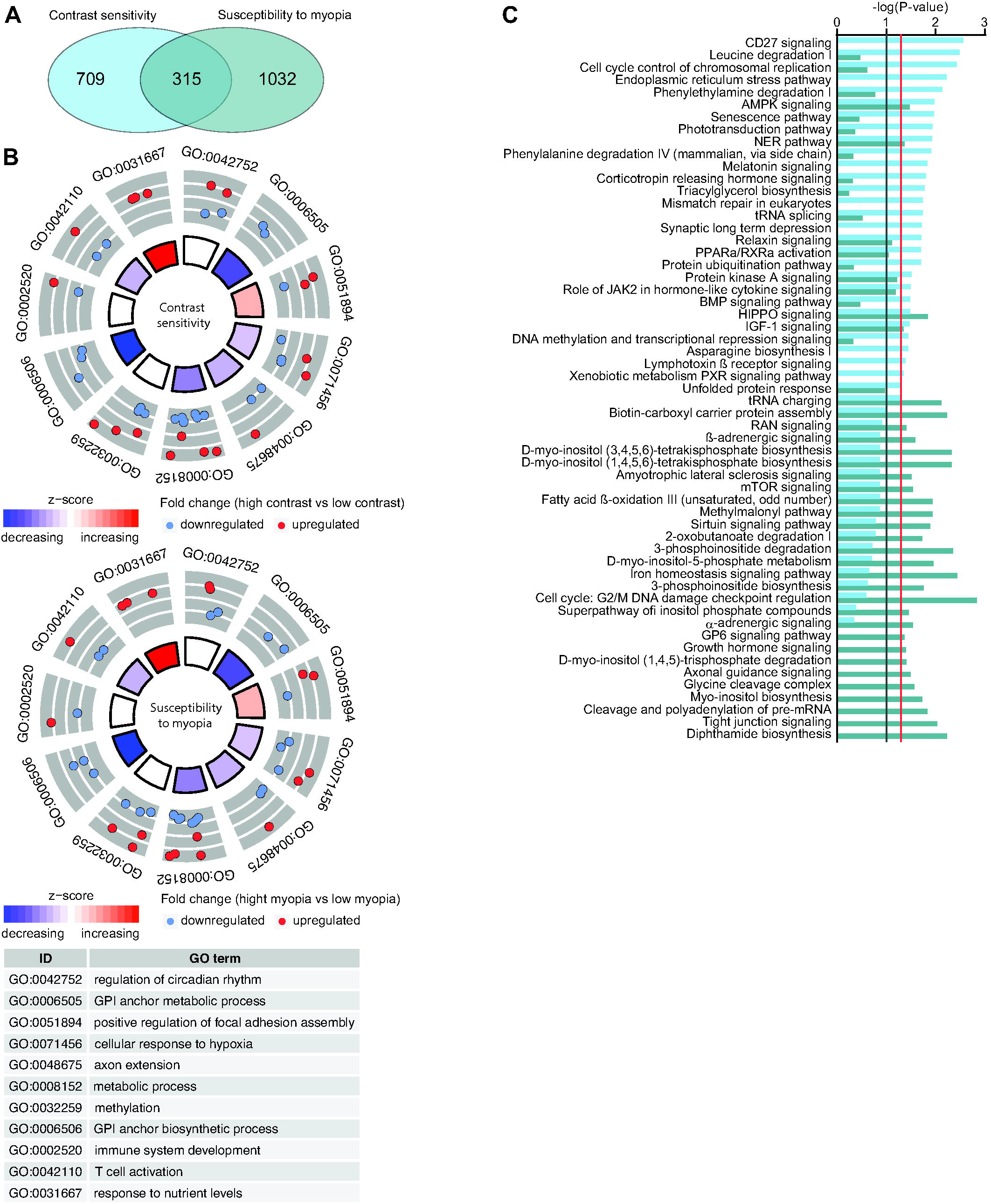
Genetic network underlying contrast perception has significant contribution to the regulation of susceptibility to form-deprivation myopia. (A) Venn diagram showing overlap between genes underlying contrast sensitivity and genes regulating susceptibility to form-deprivation myopia. (B) Top 11 biological processes affected by the genes associated with both contrast perception (top panel) and susceptibility to form-deprivation myopia in mice (bottom panel). Outer circle shows gene ontology IDs for the biological processes; middle circle shows up- or down-regulated genes in animals with high contrast sensitivity versus animals with low contrast sensitivity (top panel), or in mice with high susceptibility to form-deprivation myopia versus mice with low susceptibility to form-deprivation myopia (bottom panel); inner circle shows activation or suppression of the corresponding biological processes, while the size of the sector corresponds to statistical significance (larger sectors correspond to smaller P-values). (C) Comparison of canonical signaling pathways involved in contrast perception and susceptibility to form-deprivation myopia. Vertical red line indicates P = 0.05. Vertical grey line indicates P = 0.1. Colors identify pathways associated with either contrast perception or susceptibility to form-deprivation myopia and correspond to the colors in the Venn diagrams (A).

### Genes involved in contrast perception are linked to human myopia and several other classes of human genetic disorders

Our data suggested that genes involved in contrast perception, baseline refractive eye development, and susceptibility to form-deprivation myopia encode a variety of signaling pathways regulating a diverse range of biological processes. To obtain an additional layer of information about the spectrum of contrast-perception-related biological processes involved in refractive eye development, we analyzed the association between contrast genes, which were found to be involved in either baseline refractive development or form-deprivation myopia development, and human genetic disorders listed in the Online Mendelian Inheritance in Man (OMIM) database. We also analyzed the overlap between these genes and the genes found to be linked to human myopia by GWAS studies. We found that 26 out of 136 genes underlying both contrast perception and baseline refractive eye development were associated with known human disorders (Figure 7A, Supplementary Table S17). The majority of these genes were linked to metabolic disorders (∼25.8%), while other genes were associated with myopia (∼22.6%), developmental disorders (∼22.6%), connective tissue disorders (∼9.7%), disorders affecting phototransduction-related signaling (∼6.5%), neurologic disorders (∼6.5%), and diseases caused by the breakdown in epigenetic regulation of gene expression (∼6.5%). Forty-nine out of 315 genes involved in both contrast perception and form-deprivation myopia development were linked to known human diseases (Figure 7B, Supplementary Table S18). The largest number of these genes were associated with myopia (39.7%). Other genes were associated with metabolic disorders (27%), developmental disorders (12.7%), diseases caused by the breakdown in synaptic transmission (11.1%), disorders affecting phototransduction-related signaling (4.8%), connective tissue disorders (1.6%), epigenetic disorders (1.6%), and diseases caused by the breakdown in DNA repair (1.6%). Cumulatively, these data suggest that the contrast-perception-related genetic network contributes to baseline refractive eye development through metabolic and developmental processes, connective tissue restructuring, phototransduction-related signaling, and epigenetic regulation. These data also suggest that the contrast-perception-related genetic network plays a prominent role in optical defocus detection and emmetropization mainly through myopia-related pathways, metabolic and developmental processes, synaptic transmission, and phototransduction-related signaling.

**Figure 7.**
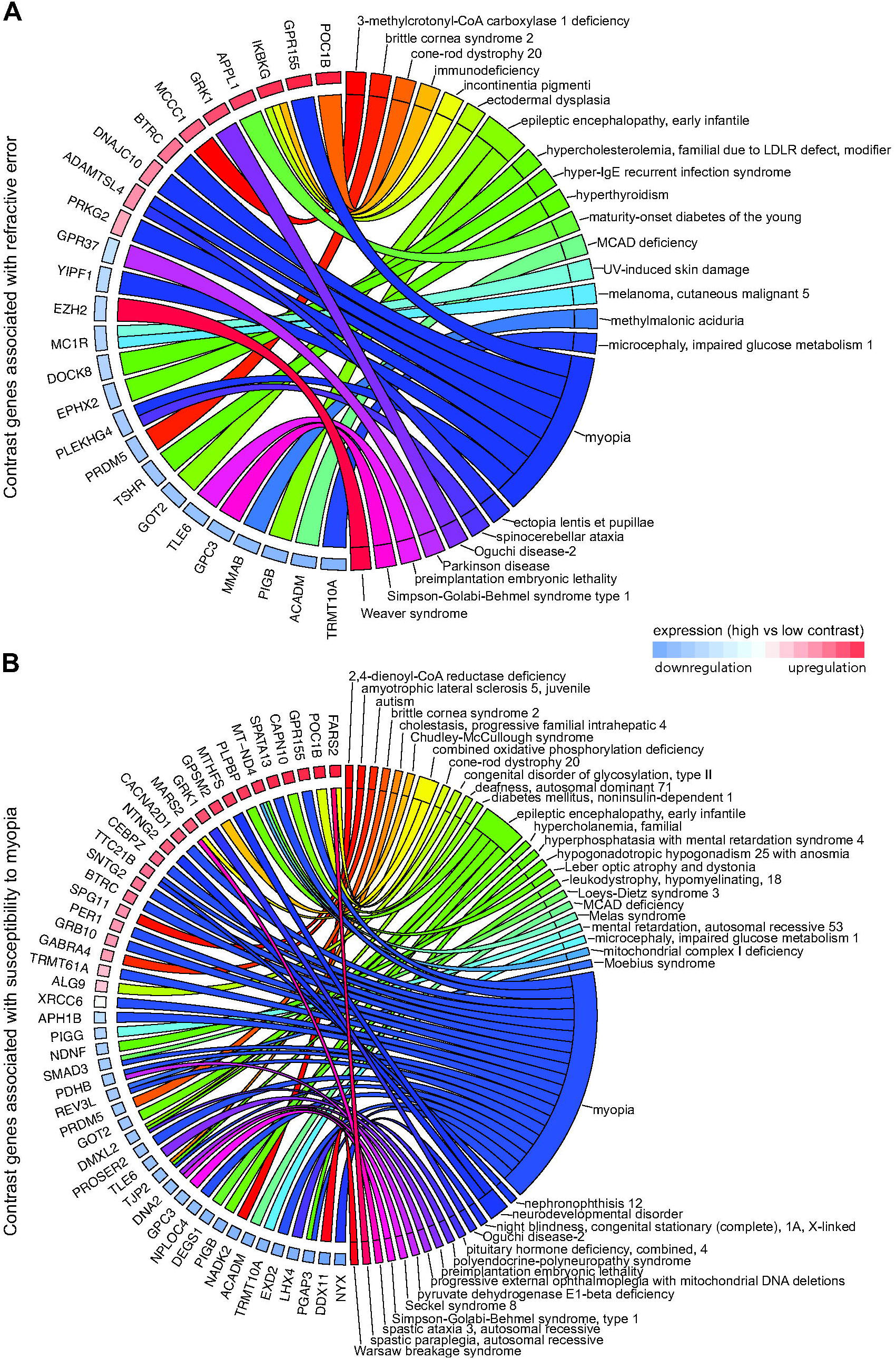
Genes underlying contrast perception are associated with diverse group of human genetic disorders. (A) Chord diagram showing genes (left semicircle) and human genetic disorders (right semicircle) associated with contrast perception and baseline refractive eye development in mice. (B) Chord diagram showing genes (left semicircle) and human genetic disorders (right semicircle) associated with contrast perception and susceptibility to form-deprivation myopia in mice. Colored bars underneath gene names show up- or down-regulation of the corresponding genes in mice with high contrast sensitivity versus mice with low contrast sensitivity.

## DISCUSSION

Several lines of evidence suggest that the process of eye emmetropization is regulated by optical defocus, which is perceived by the retina using luminance contrast and longitudinal chromatic aberrations (8, 64). Our data support this hypothesis and provide experimental data, which suggest that the genetic network and signaling pathways subserving perception of contrast by the retina play an important role in emmetropization. Although our data indicate that the signaling pathways underlying perception of contrast contribute to both baseline refractive eye development and the optical-defocus-driven emmetropization process, contrast perception plays a much more prominent role in optical defocus detection and emmetropization than in baseline refractive development. We did not find a significant correlation between contrast sensitivity and baseline refractive errors, whereas sensitivity to contrast strongly correlated with susceptibility to form-deprivation myopia. This points to the important role of contrast perception in optical defocus detection.

We found that contrast perception is strongly dependent on the signaling pathways involved in axonogenesis, synaptic signaling, cell-cell communication, and regulation of circadian rhythms. However, further analysis of the biological functions and signaling pathways subserving contrast perception revealed that the contribution of contrast-related pathways to baseline refractive eye development and optical-defocus-regulated eye emmetropization is different. While contrast-related pathways involved in baseline refractive development were primarily related to DNA methylation, histone methylation and phototransduction, contrast-related pathways underlying optical defocus detection and emmetropization were primarily involved in circadian rhythms, response to hypoxia, metabolism and synaptic transmission.

DNA methylation and phototransduction were previously implicated in refractive eye development. For example, genome-wide methylation status was linked to myopia development in humans (116, 117). In-utero epigenetic factors were found to be associated with refractive status in young children, and grandmothers’ smoking causing epigenetic modifications of the genome was shown to be linked to less myopic refractive errors in children (118–122). Light-induced signaling and the phototransduction pathway were also implicated in refractive eye development (12,113,114,123).

The finding that contrast perception is dependent on synaptic transmission is supported by previous reports that detection of contrast in the retina is organized as ON/OFF receptive fields, with a prominent role played by horizontal and amacrine cells which provide lateral inhibition via synaptic contacts (83, 84). Signaling pathways underlying response to hypoxia and metabolism were also implicated in myopia development (112, 114). However, we found that genes associated with the signaling pathways involved in the regulation of circadian rhythms were particularly overrepresented within the genetic network that controls contrast perception underlying defocus detection and emmetropization. This finding is especially intriguing because several studies found a link between circadian rhythms and refractive eye development (8, 124). Nickla et al. (125, 126) discovered that the impact of optical defocus on refractive eye development was strongly dependent on the time of day and was associated with the endogenous rhythms in choroidal thickness and eye growth. It was also found that the effect of anti-myopia drugs quinpirole and pirenzepine on myopia development depends on the time of day (127). Moreover, it was observed that contrast sensitivity strongly depends on the level of oxygen and glucose in the retina, both of which are under circadian control (128). In line with this evidence, it was reported that optical defocus alters the expression of several genes encoding endogenous eye clock (129) and that targeted retina-specific disruption of the clock gene *Bmal1* induces myopia-like phenotype in mice (130). This evidence and our results suggest that the link between circadian rhythms and sensitivity of the eye to optical defocus, hence susceptibility to optical-defocus-induced myopia, may be explained by the strong dependence of contrast perception on the retinal circadian clock signaling.

Analysis of the specific genes encoding components of the contrast-related signaling pathways led to several important findings. We found that 44% of contrast-related genes involved in the development of form-deprivation myopia in mice were linked to human myopia, while only 27% of contrast-related genes involved in baseline refractive eye development were associated with human myopia (Supplementary Tables S17 and S18). This suggests that the relative majority of genes causing human myopia are associated with pathways responsible for the processing of optical defocus.

Several of these genes deserve special attention. One of the genes, *APH1B*, encodes a critical component of the gamma-secretase complex, which is known to play an important role in the processing of amyloid beta (A4) precursor protein (APP) (131–133). APP interacts with its homologs APLP1 and APLP2 to form a presynaptic complex in neuronal axons, which plays a critical role in synaptic transmission (134–139). Importantly, *APLP2* was found to play an important role in gene-environment interaction underlying the development of childhood myopia (7). Another gene, *CACNA2D1*, which encodes a subunit of calcium voltage-gated channels mediating the influx of calcium ions into the cell upon membrane polarization, also plays an important role in synaptic transmission in the retina (140–143). Two genes encoding components of ubiquitin-protein-ligase complex, beta-transducin repeat containing E3 ubiquitin protein ligase (*BTRC*) and ubiquitin recognition factor NPL4 homolog (*NPLOC4*), confirm the importance of the protein ubiquitination pathway in myopia development identified by several studies (12,112–114,144–148). Very little is known about the function of the tight junction protein 2 encoded by the *TJP2* gene, which is primarily expressed in the inner nuclear layer of the retina (149); however, our findings and recent studies point to a potentially important role of *TJP2* in refractive eye development (113,150,151). Another interesting gene, which we found to be linked to both contrast perception and susceptibility to form-deprivation myopia, is *PER1*. The *PER1* gene encodes period circadian regulator 1 protein, which plays a critical role in the regulation of circadian rhythms (152–154). *PER1* is expressed in the inner nuclear layer of the retina harboring amacrine cells (153), which were implicated in optical defocus detection and myopia development (89–99). Interestingly *PER1* expression is regulated by the EGR1 transcription factor (also known as ZENK) and vasoactive intestinal polypeptide (VIP) (155, 156). The *EGR1* gene is expressed in the amacrine cell of the retina and was shown to respond to optical defocus in a sign-of-defocus sensitive manner (92), while VIP is the principal neurotransmitter of the VIPergic amacrine cells of the retina, which were shown to be involved in myopia development (89,157,158). Moreover, *PER1* expression is controlled by the hypoxia signaling pathway (159), which was shown to play a critical role in the signaling cascade underlying the eye’s response to optical defocus (112, 114). Thus, our data suggest an intriguing interaction between contrast perception, the retinal circadian clock pathway and the signaling cascade underlying optical defocus detection.

We also found a large number of contrast-related genes, which were not previously implicated in refractive eye development or myopia, but were found to be linked to a multitude of other human genetic disorders. Analysis of these genes also provides additional insights into biological processes involved in contrast perception and refractive eye development.

For example, several such genes associated with both contrast perception and baseline refractive development point to the involvement of several seemingly unrelated biological processes in baseline refractive eye development. The causal gene for a congenital form of cone-rod dystrophy *POC1B* and the gene causing Oguchi disease *GRK1* are involved in the functioning of photoreceptor synapses and light-dependent deactivation of rhodopsin respectively (160–162), which suggests that photoreceptor-related signaling is involved in baseline refractive eye development. The involvement of *TSHR* gene encoding thyroid stimulating hormone receptor, which causes hyperthyroidism (163), points to the important role of the thyroid-stimulating hormone signaling pathway in baseline refractive eye development. A causal gene for Parkinson disease *GPR37* implicates dopamine signaling in baseline refractive eye development (164). Finally, it was shown that a histone methyltransferase encoded by the causal gene for Weaver syndrome *EZH2* interacts the polycomb repressive complex 2 (PRC2) and directly controls DNA methylation, implicating epigenetic regulation of gene expression in baseline refractive eye development (165, 166).

On the contrary, analysis of the genes associated with both contrast perception and susceptibility to form-deprivation myopia implicates synaptic transmission and retinal ON/OFF signaling pathways in optical defocus detection and emmetropization. For example, the causal gene for juvenile amyotrophic lateral sclerosis and Kjellin syndrome *SPG11*, *GABRA4* causing autism, *DMXL2* linked to an autosomal dominant form of deafness and early infantile epileptic encephalopathy, as well as *NTNG2* associated with a neurodevelopmental disorder are all involved in synapse function and synaptic transmission (167–177). The causal gene for Chudley-McCullough syndrome *GPSM2*, *POC1B* linked to a congenital form of cone-rod dystrophy, *GRK1* associated with a congenital stationary night blindness are involved in photoreceptor functioning (160-162,178,179), while *NYX* causing a congenital form of stationary night blindness and *LHX4* linked to congenital pituitary hormone deficiency are involved in the communication between photoreceptors and cone bipolar cells (87,180–187). Considering that communication between photoreceptors and bipolar cells plays an important role in the organization of ON/OFF receptive fields (83, 84), these two groups of genes point to the important role of ON/OFF signaling pathways in contrast perception and optical defocus detection.

In conclusion, our results provide evidence that the retinal genetic network subserving contrast perception plays an important role in optical defocus detection and emmetropization. Our results reveal that contrast-related pathways involved in baseline refractive eye development are primarily related to DNA methylation, histone methylation, phototransduction, photoreceptor-bipolar cell signaling, thyroid-stimulating hormone signaling, dopamine signaling and epigenetic regulation of gene expression. Contrast-related pathways underlying optical defocus detection and emmetropization are primarily involved in retinal ON/OFF signaling pathways, synaptic transmission, circadian rhythms, response to hypoxia and metabolism. Our results suggest an interaction between contrast perception, the retinal circadian clock pathway and the signaling cascade underlying optical defocus detection. We note that the link between circadian rhythms and sensitivity of the eye to optical defocus may be explained by the strong dependence of contrast perception on the retinal circadian clock signaling. This study also suggests that the relative majority of genes causing common human myopia are involved in the processing of optical defocus, i.e. gene-environment interaction.

## ACKNOWLEDGMENTS

This work was supported by the National Institutes of Health grants R01EY023839 (AVT), P30EY019007 (Core Support for Vision Research received by the Department of Ophthalmology, Columbia University), and Research to Prevent Blindness (Unrestricted funds received by the Department of Ophthalmology, Columbia University). The funders had no role in study design, data collection and analysis, decision to publish, or preparation of the manuscript.

## AUTHOR CONTRIBUTIONS

TVT and AVT conceptualized the study, analyzed refractive eye development, susceptibility to form-deprivation myopia and contrast sensitivity in mice, performed RNA-seq, and analyzed the data. AVT supervised the entire study, analyzed and validated data, and wrote the original draft of the manuscript. All authors read, edited, and approved the final version of the manuscript.

## METHODS

### Mice

Mice comprising collaborative cross (129S1/svlmj, A/J, C57BL/6J, CAST/EiJ, NOD/ShiLtJ, NZO/HlLtJ, PWK/PhJ, and WSB/EiJ mice) were obtained from the Jackson Laboratory (Bar Harbor, ME) and were maintained as an in-house breeding colony. Food and water were provided ad libitum. All procedures adhered to the Association for Research in Vision and Ophthalmology (ARVO) statement on the use of animals in ophthalmic and vision research and were approved by the Columbia University Institutional Animal Care and Use Committee. Animals were anesthetized via intraperitoneal injection of ketamine (90 mg/kg) and xylazine (10 mg/kg) and were euthanized using CO_2_ followed by cervical dislocation.

### Analysis of contrast sensitivity

Contrast sensitivity was measured at P40 using a virtual optomotor system (Mouse OptoMotry System, Cerebral Mechanics), as previously described (188). Briefly, the animal to be tested was placed on a platform surrounded by four computer screens displaying a virtual cylinder comprising a vertical sine wave grating in 3D coordinate space. The OptoMotry software controlled the speed of rotation, direction of rotation, the frequency of the grating and its contrast. Contrast sensitivity was measured at six spatial frequencies, i.e. 0.031, 0.064, 0.092, 0.103, 0.192, and 0.272 cycles/degree (cpd) using the staircase procedure. The contrast was systematically decreased from 100% using the staircase procedure until the minimum contrast capable of eliciting a response (contrast sensitivity) was determined. The staircase procedure was such that 3 correct answers in a row advanced it to a lower contrast, while 1 wrong answer returned it to a higher contrast. The contrast sensitivity at each frequency was calculated as a reciprocal of the contrast threshold, which was calculated as a Michelson contrast from the screen luminances

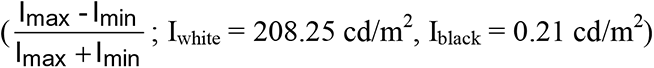

### Analysis of refractive state of the eye

We measured baseline refractive errors in both left and right eyes on alert animals at P40 using an automated eccentric infrared photorefractor as previously described (189, 190). The animal to be refracted was immobilized using a restraining platform, and each eye was refracted along the optical axis in dim room light (< 1 lux), 20-30 min. after instilling 1% tropicamide ophthalmic solution (Alcon Laboratories) to ensure mydriasis and cycloplegia. The measurements were automatically acquired by the photorefractor every 16 msec. Each successful measurement series (i.e., Purkinje image in the center of the pupil and stable refractive error for at least 5 sec.) was marked by a green LED flash, which was registered by the photorefractor software. Five independent measurement series were taken for each eye. Sixty individual measurements from each series, immediately preceding the green LED flash, were combined, and a total of 300 measurements (60 measurements x 5 series = 300 measurements) were collected for each eye. Data for the left and right eyes were combined (600 measurements total) to calculate the mean refractive error and standard deviation for each animal.

### Analysis of susceptibility to form-deprivation myopia

We measured the extent of myopia induced by diffuser-imposed retinal image degradation (visual form deprivation). Visual form deprivation was induced in one of the eyes by applying plastic diffusers and refractive development of the treated eye was compared to that of the contralateral eye, which was not treated with a diffuser, as previously described (50, 191). Diffusers represented low-pass optical filters, which degraded the image projected onto the retina by removing high spatial frequency details. Hemispherical plastic diffusers were made from zero power rigid contact lenses manufactured from OP3 plastic (diameter = 7.0 mm, base curve = 7.0 mm; Lens.com) and Bangerter occlusion foils (Precision Vision). Diffusers were inserted into 3D-printed plastic frames (Proto Labs). On the first day of the experiment (P24), animals were anesthetized via intraperitoneal injection of ketamine and xylazine, and frames with diffusers were attached to the skin surrounding the eye with six stitches using size 5-0 ETHILON™ microsurgical sutures (Ethicon) and reinforced with Vetbond™ glue (3M Animal Care Products) (the contralateral eye served as a control). Toenails were covered with adhesive tape to prevent mice from removing the diffusers. Animals recovered on a warming pad and were then housed under low-intensity constant light in transparent plastic cages for the duration of the experiment as previously described (50, 191). Following 21 days of visual form deprivation (from P24 through P45), diffusers were removed and the refractive state of both treated and control eyes was assessed using an automated infrared photorefractor as previously described (189, 190). The interocular difference in refractive error between the treated and contralateral control eye was used as a measure of induced myopia.

### RNA extraction and RNA-seq

Animals were euthanized following an IACUC-approved protocol. Eyes were enucleated, the retinae were dissected from the enucleated eyes. The retinae were washed in RNAlater (Thermo Fisher Scientific) for 5 min., frozen in liquid nitrogen, and stored at −80°C. To isolate RNA, tissue samples were homogenized at 4°C in a lysis buffer using Bead Ruptor 24 tissue homogenizer (Omni). Total RNA was extracted from each tissue sample using miRNAeasy mini kit (QIAGEN) following the manufacturer’s protocol. The integrity of RNA was confirmed by analyzing 260/280 nm ratios (Ratio_260/280_ = 2.11-2.13) on a Nanodrop (Thermo Scientific) and the RNA Integrity Number (RIN = 9.0-10.0) using Agilent Bioanalyzer. Illumina sequencing libraries were constructed from 1 μg of total RNA using the TruSeq Stranded Total RNA LT kit with the Ribo-Zero Gold ribosomal RNA depletion module (Illumina). Each library contained a specific index (barcode) and were pooled at equal concentrations using the randomized complete block (RCB) experimental design before sequencing on Illumina HiSeq 2500 sequencing system. The number of libraries per multiplexed sample was adjusted to ensure sequencing depth of ∼70 million reads per library (paired-end, 2 x 100 bp). The actual sequencing depth was 76,773,554 ± 7,832,271 with read quality score 34.5 ± 0.4.

### Post-sequencing RNA-seq data validation and analysis

The FASTQ raw data files generated by the Illumina sequencing system were imported into Partek Flow software package (Partek), libraries were separated based on their barcodes, adapters were trimmed and remaining sequences were subjected to pre-alignment quality control using the Partek Flow pre-alignment QA/QC module. After the assessment of various quality metrics, bases with the quality score < 34 were removed (≤ 5 bases) from each end. Sequencing reads were then mapped to the mouse reference genome Genome Reference Consortium Mouse Build 38 (GRCm38/mm10, NCBI) using the STAR aligner resulting in 95.0 ± 0.4% mapped reads per library, which covered 35.4 ± 1.0% of the genome. Aligned reads were quantified to transcriptome using the Partek E/M annotation model and the NCBI’s RefSeq annotation file to determine read counts per gene/genomic region. The generated read counts were normalized by the total read count and subjected to the analysis of variance (ANOVA) to detect genes whose expression correlates with either refractive error, susceptibility to myopia or contrast sensitivity. Differentially expressed transcripts were identified using a P-value threshold of 0.05 adjusted for genome-wide statistical significance using Storey’s q-value algorithm (192). To identify sets of genes with coordinate expression, differentially expressed transcripts were clustered using the Partek Flow hierarchical clustering module, using average linkage for the cluster distance metric and Euclidean distance metric to determine the distance between data points. Each RNA-seq sample was analyzed as a biological replicate, thus, resulting in three biological replicates per strain.

### Gene ontology analysis and identification of canonical signaling pathways

To identify biological functions (gene ontology categories), which were significantly associated with the genes whose expression correlated with baseline refractive errors, susceptibility to myopia or contrast sensitivity, we used the database for annotation, visualization and integrated discovery (DAVID) version 6.8 (193) and GOplot R package (194). DAVID uses a powerful gene-enrichment algorithm and DAVID Gene Concept database to identify biological functions (gene ontology categories) affected by differential genes, while GOplot integrates gene ontology information with gene expression information and predicts the effects of gene expression changes on biological processes. DAVID uses a modified Fisher’s exact test (EASE score) with a P-value threshold of 0.05 to estimate the statistical significance of enrichment for specific gene ontology categories. The IPA Pathways Activity Analysis module (QIAGEN) was used to identify canonical pathways encoded by the genes involved in baseline refractive eye development, susceptibility to myopia or contrast perception, and to predict the effects of gene expression differences in different mouse strains on the pathways. The activation z-score was employed in the Pathways Activity Analysis module to predict activation or suppression of the canonical pathways. The z-score algorithm is designed to reduce the chance that random data will generate significant predictions. The z-score provides an estimate of statistical quantity of change for each pathway found to be statistically significantly affected by the changes in gene expression. The significance values for the canonical pathways were calculated by the right-tailed Fisher’s exact test. The significance indicates the probability of association of molecules from a dataset with the canonical pathway by random chance alone. The Pathways Activity Analysis module determines if canonical pathways, including functional end-points, are activated or suppressed based on the gene expression data in a dataset. Once statistically significant canonical pathways were identified, we subjected the datasets to the Core Functional Analysis in IPA to compare the pathways and identify key similarities and differences in the canonical pathways underlying baseline refractive development, susceptibility to myopia and contrast sensitivity.

### Identification of genes linked to human myopia and other human genetic disorders

All mouse genes, which were found to be associated with baseline refractive errors, susceptibility to myopia or contrast sensitivity were analyzed for their association with human genetic disorders. To identify genes linked to human myopia, we compared the genes that we found in mice with a list of genes located within human myopia QTLs. We first compiled a list of all SNPs or markers exhibiting a statistically significant association with myopia in the human linkage or genome-wide association studies (GWAS) using the Online Mendelian Inheritance in Man (OMIM) (McKusick-Nathans Institute of Genetic Medicine, Johns Hopkins University) and NHGRI-EBI GWAS Catalog (195) databases. The LDlink’s LDmatrix tool (National Cancer Institute) was used to identify SNPs in linkage disequilibrium and identify overlapping chromosomal loci. We then used the UCSC Table Browser to extract all genes located within critical chromosomal regions identified by the human linkage studies or within 200 kb (±200 kb) of the SNPs found by GWAS. The list of genes located within human QTLs was compared with the list of genes that we found in mice using Partek Genomics Suite (Partek). To identify genes associated with human genetic disorders unrelated to myopia, we screened mouse genes that we found in this study against the Online Mendelian Inheritance in Man (OMIM) (McKusick-Nathans Institute of Genetic Medicine, Johns Hopkins University) database.

### Statistical data analysis and data graphing

Statistical analyses of the RNA-seq data were performed using statistical modules integrated into Partek Flow (Partek), DAVID (193) and IPA (QIAGEN) software packages and described in the corresponding sections. Other statistical analyses were performed using the STATISTICA software package (StatSoft). The statistical significance of the overlaps between gene datasets was estimated using probabilities associated with the hypergeometric distribution as implemented in the Bioconductor R software package GeneOverlap and associated functions. Data graphing and visualization was performed using Partek Flow (Partek) and IPA (QIAGEN) graphing and visualization modules, as well as Prism 8 for Windows (GraphPad) and GOplot R package (194).

## SUPPLEMENTAL INFORMATION

**Table S1.** Refractive errors in mice comprising collaborative cross (P40, diopters)

**Table S2.** Form-deprivation myopia in mice comprising collaborative cross (deprived eye versus control eye, diopters)

**Table S3.** Contrast sensitivity in mice comprising collaborative cross

**Table S4.** Genes whose expression correlates with refractive error in mice comprising collaborative cross (FC = fold change, hyperopia versus myopia)

**Table S5.** Genes whose expression correlates with susceptibility to myopia in mice comprising collaborative cross (FC = fold change, high versus low myopia)

**Table S6.** Genes whose expression correlates with contrast sensitivity in mice comprising collaborative cross (FC = fold change, high versus low contrast sensitivity)

**Table S7.** Gene ontology categories associated with genes whose expression correlates with refractive error in mice comprising collaborative cross (BP, biological process; CC, cellular component; MF, molecular function)

**Table S8.** Gene ontology categories associated with genes whose expression correlates with susceptibility to myopia in mice comprising collaborative cross (BP, biological process; CC, cellular component; MF, molecular function)

**Table S9.** Gene ontology categories associated with genes whose expression correlates with contrast sensitivity in mice comprising collaborative cross (BP, biological process; CC, cellular component; MF, molecular function)

**Table S10.** Canonical pathways associated with genes whose expression correlates with refractive error in mice comprising collaborative cross

**Table S11.** Canonical pathways associated with genes whose expression correlates with susceptibility to myopia in mice comprising collaborative cross

**Table S12.** Canonical pathways associated with genes whose expression correlates with contrast sensitivity in mice comprising collaborative cross

**Table S13.** Genes whose expression correlates with both contrast sensitivity and refractive error in mice comprising collaborative cross (FC = fold change, contrast sensitivity = high versus low contrast sensitivity, RE = hyperopia versus myopia)

**Table S14.** Genes whose expression correlates with both contrast sensitivity and susceptibility to myopia in mice comprising collaborative cross (FC = fold change, contrast sensitivity = high versus low contrast sensitivity, myopia = high versus low myopia)

**Table S15.** Gene ontology categories associated with genes whose expression correlates with both contrast sensitivity and refractive error in mice comprising collaborative cross (BP, biological process; CC, cellular component; MF, molecular function)

**Table S16.** Gene ontology categories associated with genes whose expression correlates with both contrast sensitivity and susceptibility to myopia in mice comprising collaborative cross (BP, biological process; CC, cellular component; MF, molecular function)

**Table S17.** Genes whose expression correlates with both contrast sensitivity and refractive error in mice comprising collaborative cross and linked to human diseases (FC = fold change, high versus low contrast sensitivity)

**Table S18.** Genes whose expression correlates with both contrast sensitivity and susceptibility to myopia in mice comprising collaborative cross and linked to human diseases (FC = fold change, high versus low contrast sensitivity)

